# Feedback inhibition of AMT1 NH_4_^+^-transporters mediated by CIPK15 kinase

**DOI:** 10.1101/2020.11.03.367383

**Authors:** Hui-Yu Chen, Yen-Ning Chen, Hung-Yu Wang, Zong-Ta Liu, Wolf B. Frommer, Cheng-Hsun Ho

## Abstract

Ammonium (NH_4_^+^), a key nitrogen form, becomes toxic when it accumulates to high levels. Ammonium transporters (AMTs) are the key transporters responsible for NH_4_^+^ uptake. AMT activity is under allosteric feedback control, mediated by phosphorylation of a threonine in the cytosolic C-terminus (CCT). However, the kinases responsible for the NH_4_^+^-triggered phosphorylation remain unknown. In this study, a functional screen identified protein kinase <u>C</u>BL-<u>I</u>nteracting <u>P</u>rotein <u>K</u>inase15 (CIPK15) as a negative regulator of AMT1;1 activity. CIPK15 was able to interact with several AMT1 paralogs at the plasma membrane. Analysis of AmTryoshka, an NH_4_^+^ transporter activity sensor for AMT1;3 in yeast, and a two-electrode-voltage-clamp (TEVC) of AMT1;1 in *Xenopus* oocytes showed that CIPK15 inhibits AMT activity. CIPK15 transcript levels increased when seedlings were exposed to elevated NH_4_^+^ levels. Notably, *cipk15* knockout mutants showed higher ^15^NH_4_^+^ uptake and accumulated higher amounts of NH_4_^+^ compared to the wild-type. Consistently, *cipk15* was hypersensitive to both NH_4_^+^ and methylammonium but not nitrate (NO_3_^−^). Taken together, our data indicate that feedback inhibition of AMT1 activity is mediated by the protein kinase CIPK15 via phosphorylation of residues in the CCT to reduce NH_4_^+^-accumulation.

## INTRODUCTION

As a key building block of nucleic acids, amino acids, and proteins, nitrogen is an essential nutrient. In plants, nitrogen supply can limit or inhibit growth, development and crop yield when below or above the optimal range. Ammonium (NH_4_^+^) is one of the main inorganic forms of nitrogen for plant nutrition. NH_4_^+^ is also an important nitrogen source for bacteria, fungi, and plants, but becomes toxic when it rises above certain levels [1–5].

Plants take up NH_4_^+^ with the help of specific transporters. AMT/MEP/Rhesus protein superfamily members function as electrogenic high-affinity NH_4_^+^ transporters [6–9]. Potassium (K^+^) channels can also mediate NH_4_^+^ uptake [10]. The *Arabidopsis* genome contains six AMT paralogs, four (AMT1;1, 1;2, 1;3, and 1;5) of which are together essential for NH_4_^+^ uptake [11, 12]. Unlike K^+^ channels, AMTs are highly selective for NH_4_^+^ and its methylated form, methylammonium (MeA) [6, 13]. In addition to their roles as transporters, AMTs can also function as receptors involved in the control of root growth and development, similar to the yeast MEP2 transceptor, which measures NH_4_^+^ concentrations to regulate pseudohyphal growth [14]. Recently, a ratiometric biosensor of NH_4_^+^ transporter activity, named AmTryoshka, which reports NH_4_^+^ transporter activity *in vivo*, was developed by inserting a cassette carrying two fluorophores into AMT1;3 [15]. Most organisms, including animals, plants, and even bacteria are sensitive to high levels of NH_4_^+^. A sole supply of nitrogen as NH_4_^+^ is typically noxious [16]. In bacteria, the existence of highly effective detoxification mechanisms may have prevented the discovery of NH_4_^+^ toxicity. The actual mechanisms of NH_4_^+^ toxicity are not understood for any organism, but several hypotheses have been proposed: (i) pH effects-uptake and assimilation of NH_4_^+^ lead to acidification of the cytosol; (ii) membrane depolarization-NH_4_^+^ uptake depolarizes the membrane, thus high levels of NH_4_^+^ uptake could affect the capability of the cell to take up other nutrients; (iii) inhibition of electron flow in plastidic and mitochondrial membranes; (iv) increased production of reactive oxygen species that damage the cells [17, 18]; and (v) replacement of potassium as an enzyme cofactor, altering the catalytic properties and/or folding of enzymes that require K^+^ [19, 20]. The increased NH_4_^+^ toxicity at low K^+^ concentration strongly supports the K^+^ replacement hypothesis [21]. Overaccumulation of ammonium can occur under various conditions, e.g., due to local placement of high doses by animals or by overfertilization. The concentrations of salts, and thus ammonium also rapidly increase during soil drying. To prevent accumulation of NH_4_^+^ becoming toxic, the activity of AMTs is tightly regulated and likely based on feedback inhibition [22, 23]. Recent reports indicate that the phosphorylation of critical threonine (T460), which is triggered by NH_4_^+^ in the cytosolic C-terminus (CCT) of AMT1;1 leads to transport inhibition via allosteric regulation in the trimeric transporter complex [24, 25]. The AMT1;1 CCT, which serves as an allosteric switch, is highly conserved among the AMT homologs found in species ranging from bacteria to higher plants. Use of this allosteric regulation mechanism of AMT1;1 for feedback control allows plants to rapidly and efficiently block the uptake of NH_4_^+^ before levels become toxic. Yet, the full circuitry leading to NH_4_^+^-dependent phosphorylation of AMTs is not fully understood. We speculate that specific kinases are activated under conditions that lead to NH_4_^+^ accumulation.

Members of the CBL-interacting protein kinases [CIPK, also named SNF1-related kinases (SnRK)], which typically function together with members of the Calcineurin B-like protein (CBL) family, are known to regulate the activities of diverse types of transporters including the plasma membrane Na^+^/H^+^ exchanger SOS1 [26], the potassium channel AKT1 [27], magnesium transport [28], the nitrate transceptor CHL1 [29], the H^+^/ATPase AHA2 [30] and several anion channels [31]. To examine whether CIPKs function as AMT regulators, we systematically screened for CIPKs able to affect AMT1;1 activity in *Xenopus* oocytes. In this study, we show that *Arabidopsis* CIPK15 acts as a negative regulator of AMT1;1 activity. CIPK15 directly interacted with AMT1;1 and inhibited AMT1;1 activity via phosphorylation of T460. This negative effect of CIPK15 on AMT1;1 activity was also observed by using an NH_4_^+^ transporter activity sensor-AmTryoshka1;3 LS-F138I, a ratiometric genetically encoded biosensor in yeast [15]. *CIPK15* transcript levels increased in response to addition of external NH_4_^+^. Notably, compared to wild-type, *cipk15* mutants showed higher ^15^NH_4_^+^ uptake and accumulated higher amounts of NH_4_^+^ and were hypersensitive to both NH_4_^+^ and MeA. Together, our data indicate that in the presence of elevated NH_4_^+^, CIPK15 inhibits AMT1;1 activity to prevent NH_4_^+^ toxicity.

## MATERIALS AND METHODS

### Plant materials and treatments

Experiments were performed with *Arabidopsis thaliana* ecotype Col-0. The knockout lines of AMT-*qko*, the quadruple AMT which carry T-DNA insertions in *AMT1;1*, *1;2*, *1;3*, and *2;1*), *cbl4* (At5g24270, SALK_113101), and *cipk19* (At5g45810, SALK_044735) have been previously described [32–34]. Plant growth conditions have also been previously described [25], and were used here with minor modifications. Arabidopsis seeds were surface sterilized and germinated on half-strength modified Murashige Skoog medium (MS), nitrogen-free salts (Phytotechlab, M407) with 5 mM KNO_3_ as the sole nitrogen source, 0.5% [w/v] sucrose, and 1% [w/v] agar, pH 5.8 [KOH] on vertical plates. For qRT-PCR, protein blots, NH_4_^+^ content, and N^15^ labeled uptake assays, seedlings (7 days after germination) were transferred to the half-strength MS medium lacking nitrogen for 2 days, and then, seedlings were transferred to half-strength MS medium supplemented with NH_4_Cl, ^15^NH_4_Cl or NO_3_^−^ according to the concentrations indicated in the respective figure legends. Roots were collected and frozen in liquid nitrogen. Seedlings were incubated in a 16/8 h light/dark period at 22°C. For NH_4_^+^ content, seedlings were collected after being starved for 2 days, or after treatment with 1 mM NH_4_Cl, and 1 mM KNO_3_ for 1 h. For N^15^ labeled uptake assays, seedlings were collected after being starved for 2 days (1 mM ^15^NH_4_Cl was used for the last 15 mins for ^15^N-labeling), or after treatment with 1mM NH_4_Cl, and 1mM KNO_3_ for 1 h (1mM ^15^NH_4_Cl was used for the last 15 mins for N^15-^labeling). ^15^N-labeled seedlings were then dried for 2-3 days at 65°C and further analyzed by Thermo Finnigan Delta plus XP IRMS (ThermoFisher Scientific). For primary root length determination, seedlings (3 days after germination) were transferred to half-strength MS medium with nitrogen-free salts, containing KNO_3_, NH_4_Cl or methylammonium (MeA) as nitrogen sources at the concentrations indicated in the figures, and grown for another one to five days. Seedlings were scanned on a flatbed scanner, and primary root length was measured using NIH ImageJ software (imagej.nih.gov).

### Characterization of T-DNA insertion mutants

Col-0, *CIPK15* (*SnRK3.1*) (At5g01810) T-DNA insertion lines *cipk15-1* (SALK_203150) and *cipk15*-2 (GK604B06) were obtained from the Arabidopsis Biological Resource Center (http://www.arabidopsis.org/abrc/). T-DNA insertions in *CIPK15* were confirmed by PCR analysis and sequencing using the T-DNA left border primer (5’-TGGTTCACATAGTGGGCCATCA) and CIPK15 F (5’-TCTTCTGGTGGTAGGACACG) and R (5’-TGGAATTCCAATGTGTCACC) primers. *H3G1* (At4g40040) was used as the loading control. *H3G1,* forward primer: 5’-AACCACTGGAGGAGTCAAGA-3’; reverse primer: 5’-CAATTAAGCACGTTCTCCTCT-3’).

### Real-Time qRT-PCR analyses

Real-time qRT-PCR was performed as previously described [29]. In brief, template cDNA samples were prepared using 4 κg of total RNA and the Improm-II reverse transcription system (Promega). Primers were designed to have a T_m_ of ~60°C and to produce PCR products of ~200−400 base pairs. Expression levels in each experiment were first normalized to the expression of *Ubiquitin10* measured in the same cDNA samples (*AMT1;1*, forward primer: 5’-ACGACATTATCAGTCGC; reverse primer: 5’-CTGTCCTGTGTAGATTAACG; *CIPK15*, forward primer: 5’-GGCTACGCATCTGACT; reverse primer: 5’-CGTGCAAGCGACTATC; *CIPK23,* forward primer: 5’ TCTTCTGGTGGTAGGACACG; reverse primer: 5’-TGGAATTCCAATGTGTCACC, and *Ubiquitin10*, forward primer: 5’-CTTCGTCAAGACTTTGACCG; reverse primer: 5’-CTTCTTAAGCATAACAGAGACGAG).

### Quantification of ammonium levels in plants

NH_4_^+^ content was determined colorimetrically at 410 nm after reaction with Nessler’s reagent [35]. In brief, 500 mg of fresh matter was added to 1 ml of deionized water and shaken for 1 h at 45°C. Samples were centrifuged at 15,000*g* for 20 min. Ammonium content was determined on 50 κl of the supernatant using 1 ml of Nessler’s reagent (Merck) and quantified by using a standard curve and expressed as κmol g^−1^ FW.

### Split-ubiquitin yeast two-hybrid assays

The split-ubiquitin yeast two-hybrid assay was as described previously [36]. In brief, ORFs of interest were cloned in frame with either the C-terminal (Cub) of TMBV4 vector or N-terminal (NubG; wild-type I-13 replaced by G) domain of ubiquitin in pDL2Nx vector, and then introduced into yeast strains AP4 and AP5 by the lithium acetate method [37]. For interaction growth assays, yeast was transformed with plasmids containing AMT1s-Cub, CIPK15-Nub, CIPK19-Nub, NubI, or NubG. Colonies were picked and cells were serially diluted four-fold and grown for 2 days in either SD-Trp Leu (control) or SD-Trp Leu His (for interaction). β-galactosidase (β-gal) activity was determined using filter assays, X-gal staining, and quantitative β-gal assays [38]. For filter assays, cells were streaked on filter paper, briefly frozen in liquid nitrogen, defrosted, and placed in Petri dishes filled with 0.5% agarose containing 35 mM β-mercaptoethanol (v/v) and 1.5 mg/mL of 5-bromo-4-chloro-3-indolyl-β-D-galactoside (Sigma). For X-gal staining, yeast co-expressing bait and prey fusions were streaked onto minimal medium lacking leucine and tryptophan and onto media plates supplemented with X-gal. For quantitative β-gal assays, cells were grown in minimal medium lacking leucine and tryptophan at 30°C overnight to OD_600_ of ~0.75, centrifuged, and washed in 1 ml Buffer Z (113 mM Na_2_HPO_4_, 40 mM NaH_2_PO_4_, 10 mM KCl, and 1 mM MgSO_4_). To perform the assay, 300 μl Buffer Z was added to the pellets and vortexed before lysing cells by 3 freeze-thaw cycles. Lysate (100 μl) was added immediately to 700 μl Buffer Z containing 0.27% β-mercaptoethanol before addition of 160 μl of 4 mg/ml 2-nitrophenyl-beta-D-galactopyranoside (ONPG) in buffer Z. The lysate was incubated at 30°C for 180 min. Reactions were stopped by adding 0.4 ml of 0.1 M Na_2_CO_3_. Samples were centrifuged, and OD_420_ of the supernatant was measured. For each prey-bait combination, five independent colonies were taken and the results were averaged.

### Split-fluorescent protein interaction assays in tobacco leaves

Potential AMT1;1 and CIPK interactions were tested in a tobacco transient expression system using a modified split-fluorescent protein assay as previously described [39]. In brief, AMT1;1 and CIPKs were PCR amplified, cloned into the Gateway entry vector pENTR/D/TOPO, and then recombined into the Gateway binary destination vectors pXNGW, pNXGW, pCXGW, and pXCGW using LR clonase (Invitrogen). Each protein was independently tagged with cCFP and nYFP at either the N or C terminus. The binary vector backbone was derived from pPZP312, which contains a single 35S cauliflower mosaic virus promoter and terminator derived from pRT100. The binary constructs were further introduced into *A. tumefaciens* strain GV3101. Cell density was adjusted with infiltration buffer to OD_600_ ~0.5. Agrobacteria harboring the Tomato Bushy Stunt Virus P19 silencing suppressor were co-infiltrated to reduce gene silencing. Aliquots (0.5 ml) of Agrobacterium cells carrying a split-fluorescent protein fusion construct and P19 constructs were mixed. A syringe was used to infiltrate the mixture into the abaxial side of *N. benthamiana* leaves. Plants were incubated in a growth chamber at 22°C, with a 16-/8-h day/night cycle for 36 to 48 h. Reconstitution of yellow fluorescent protein (YFP) fluorescence, chlorophyll, and bright field images in the transformed *N*. *benthamiana* leaves were recorded using confocal fluorescence microscopy (LSM780; Carl Zeiss).

### Extraction of membrane fractions and protein gel blot analyses

For membrane preparation, roots or oocytes were ground in liquid nitrogen and resuspended in buffer containing 250 mM Tris-Cl, pH 8.5, 290 mM sucrose, 25 mM EDTA, 5 mM β-mercaptoethanol, 2 mM DTT, 1 mM phenylmethylsulfonyl fluoride (PMSF), 0.53 mM Complete Protease Inhibitor Cocktail (Sigma-Aldrich), and 0.53 mM PhosStop Phosphatase Inhibitor Cocktail (Roche Applied Science). After centrifugation at 10,000*g* for 15 min, supernatants were filtered through Miracloth (Calbiochem) and recentrifuged at 100,000*g* for 45 min. The sediment containing the microsomes was resuspended in storage buffer [400 mM mannitol, 10% glycerol, 6 mM MES/Tris, pH 8, 4 mM DTT, 2 mM PMSF, and 13 mM phosphatase inhibitor cocktails 1 and 2 (Sigma-Aldrich)]. Proteins were denatured in loading buffer (62.5 mM Tris-HCl, pH 6.8, 10% [v/v] glycerol, 2% [w/v] SDS, 0.01% [w/v] bromophenol blue, and 1% PMSF), incubated at 37°C for 30 min with or without 2.5% [v/v] β-mercaptoethanol at 0°C, and then electrophoresed in 10% SDS polyacrylamide gels (Invitrogen) and transferred to polyvinylidene fluoride membranes. Proteins were detected using the anti-AMT1;1 antibody or the anti-P-AMT1;1T460 antibody [25]. Blots were developed using an ECL Advance Western Blotting Detection Kit (Amersham). Protein and phosphorylation levels were measured using ImageJ software.

### Two-electrode voltage clamp of AMT1;1 in *Xenopus* oocytes

Two-electrode voltage clamp (TEVC) measurements were performed in *Xenopus* oocyte*s* as previously described [40]. In brief, ORFs of AMT1;1, CBL1, the constitutively active CIPK19-CA (Thr186 to Asp), and the 10 CIPKs (CIPK2, 3, 8, 9, 10, 15, 20, 23, 24, 26 and CBL1, 4, kind gift from Jörg Kudla, Münster, Germany) in two pools of 5 in Gateway pDONR221 donor vector were further cloned into pOO2-GW via LR reactions of basic of Gateway Cloning Protocols (https://www.thermofisher.com/tw/en/home/life-science/cloning/gateway-cloning/protocols.html) using LR Clonase II enzyme (Invitrogen). For *in vitro* transcription, pOO2GW plasmids were linearized with *Mlu*I or another suitable restriction enzyme. Capped cRNA was *in vitro* transcribed by SP6 RNA polymerase using mMESSAGE mMACHINE kits (Ambion, Austin, TX). *Xenopus laevis* oocytes were obtained from Ecocyte Bio Science (Austin, TX). Oocytes were injected with distilled water (50 nl as control) or cRNA from AMT1;1, CIPKs, CBL1, CBL4, or CIPK19-CA (0.5 ng to 50 ng of cRNA as indicated in figure legends in 50 nl) using a robotic injector (Multi Channel Systems, Reutlingen, Germany) [41, 42]. Cells were kept at 16°C for 2-4 days in ND96 buffer containing 96 mM NaCl, 2 mM KCl, 1.8 mM CaCl_2_, 1 mM MgCl_2_, 5 mM HEPES, pH 7.4, and gentamycin (50 μg/μl). Electrophysiological analyses were typically performed 2-3 days after cRNA injection as previously described [40]. Typical resting potentials were about −40 mV. For current (I)-voltage (V) curves, measurements were recorded from oocytes that were first clamped at −40 mV followed by a step protocol to determine voltage dependence (−20 to −200 mV for 300 ms; in −20 mV increments). The current-voltage relationships were measured by the TEVC Roboocyte system (Multi Channel Systems) [41, 43].

### Fluorimetric analyses of AmTryoshka LS-F138I with CIPKs in yeast

Fluorimetric analyses were performed in yeast as previously described [15]. In brief, CIPK15, CIPK19, and CIPK15m (K41N, an inactive form) were introduced into yeast expressing AmTryoshka1;3 LS-F138I. Vector only was used as the control. Cells were analyzed in 96-well, flat-bottom plates (Greiner Bio-One, Germany). Steady-state fluorescence was recorded using a fluorescence microplate reader (Infinite, M1000 pro, Tecan, Switzerland) in bottom-reading mode using 7.5 nm bandwidth and a gain of 100. The fluorescence emission spectra (*λ*_exc_ 440 or 485 nm; *λ*_em_ 510 or 570 nm) were background subtracted using yeast cells expressing a non-florescent vector control.

## RESULTS

### CIPK15 can block the activity of AMT1;1 and 1;3

To identify members of the CIPK family that can modulate AMT1;1 activity, two sets of mixtures of five CIPKs grouped according to their phylogenetic relationships [44] were co-expressed with AMT1;1 in *Xenopus* oocytes and AMT1;1 activity was recorded by two-electrode voltage clamping (TEVC) of *Xenopus* oocytes (Fig. S1). NH_4_^+^-induced inward currents were completely blocked by a mixture of CIPK2, 10, 15, 20 and 26, while the combination of CIPK3, 8, 9, 23 and 24 had no major impact on AMT1;1 activity (Fig. S1). The mixture of CIPK cRNAs was deconvoluted by testing individual CIPKs. During the first round of deconvolution, oocytes co-injected with equal amounts of *AMT1;1* and *CIPK15* cRNA showed strong inhibition of NH_4_^+^-induced inward currents of AMT1;1 (Fig. 1a-b). We therefore focused on CIPK15. Full inhibition of detectable AMT1;1-mediated NH_4_^+^-induced inward currents was also obtained when tenfold lower amounts of *CIPK15* cRNA (0.5 ng) were co-injected with *AMT1;1* (5 ng), a reduction of inward currents to below the detection level (Fig. 1c). Even at low *CIPK15* cRNA levels, CBL1 did not lead to detectable activation of AMT1;1 activity (Fig. S2). The inhibition of AMT1;1 activity was not due to effects of CIPK15 co-expression on AMT1;1 levels as shown by protein gel blots (Fig. S3). Because we focused on CIPK15, we cannot exclude the possibility that other CIPKs may also affect AMT activity, in particular when co-expressed with CBLs. A ratiometric fluorescence biosensor for AMT1 activity, named AmTryoshka, which reports NH_4_^+^ transporter activity *in vivo* was previously engineered by inserting a cassette carrying two fluorophores into AMT1;3 [15]. AmTryoshka1;3 LS-F138I sensor shows a reduction in the 510 to 570 nm emission ratio when it is challenged with NH_4_^+^. Since the phosphorylation site T460 in AMT1;1 is conserved in AMTs (Fig. S4), we tested whether CIPK15 affects AMT1;3 activity by measuring the response of the ratiometric AMT activity sensor AmTryoshka1;3 LS-F138I in yeast (Fig. 2). Addition of NH_4_^+^ to yeast cells expressing the sensor led to a reduction in the relative 510 to 570 nm emission ratio. CIPK15, but not its kinase inactive form (CIPK15m) blocked NH_4_^+^-induced AmTryoshka LS-F138I responses (Fig. 2). Unlike CIPK15, CIPK19, which shares 52% identity with CIPK15, did not impair NH_4_^+^-triggered AmTryishka1;3 LS-F138I response in yeast (Fig. S5). These data show that CIPK15 can exert its effect in different heterologous systems and can specifically inhibit both AMT1;1 and AMT1;3.

**Fig. 1.**
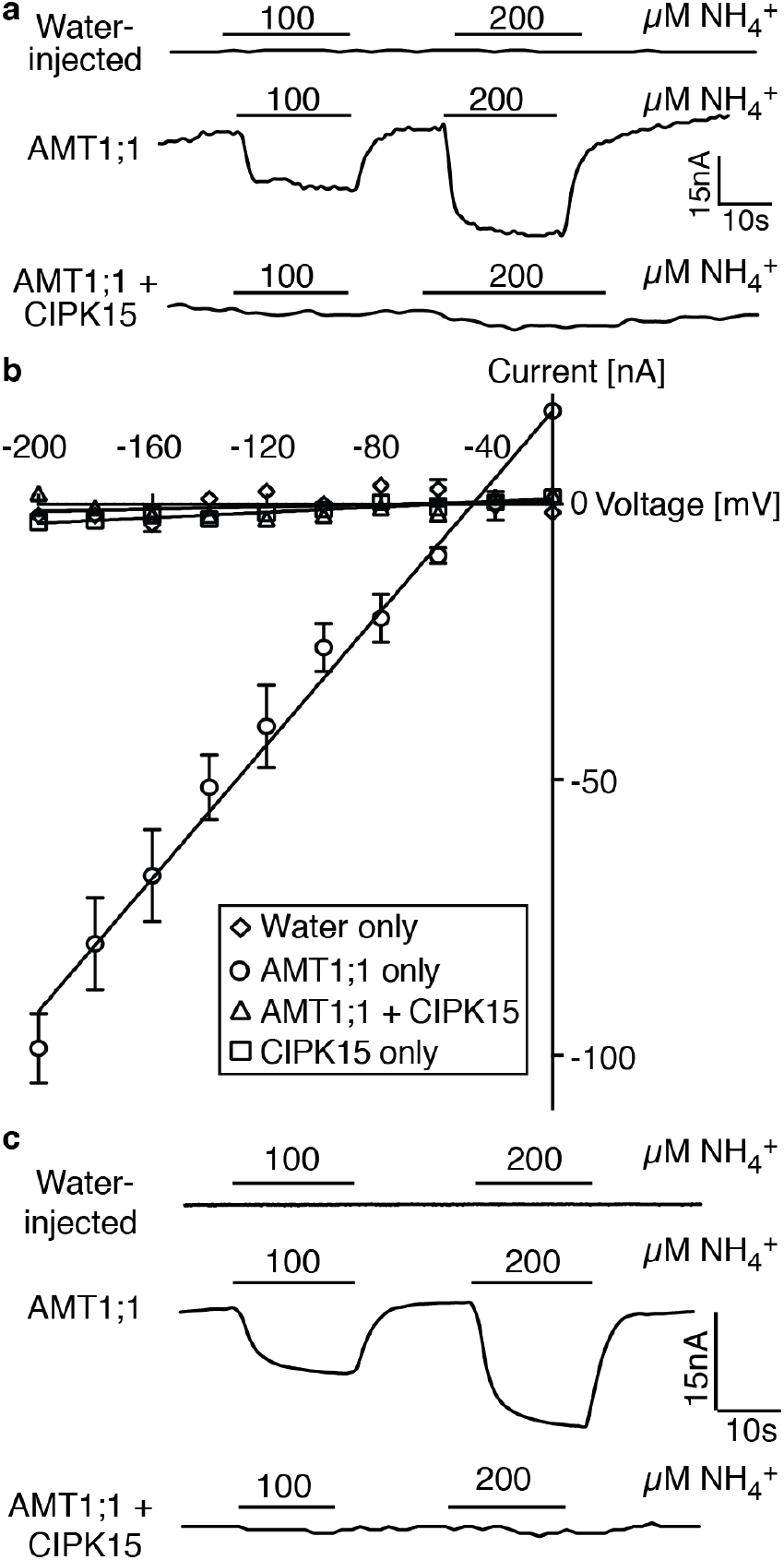
CIPK15 inhibits AMT1 activity in *Xenopus* oocytes. (**a**-**b**) Co-expression of CIPK15 inhibited NH_4_^+^-triggered inward currents of AMT1;1 in *Xenopus* oocytes. Oocytes were injected with water only, 50 ng cRNA of AMT1;1 only, 50 ng AMT1;1 + 50 ng CIPK15, or 50 ng CIPK15 only, and perfused with NH_4_Cl at the indicated concentrations (**a**) or (**b**) 0.2 mM for current recordings (**a**) and IV curve (**b**). Oocytes were voltage clamped at (**a**) −120 mV or (**b**) −40 mV and stepped in −20-mV increments between −20 and −200 mV for 300 ms. (**b**) Currents (nA) were background subtracted (difference between currents at +300 ms in the cRNA-injected AMT1;1 only/AMT1;1 + CIPK15/CIPK15 only and water-injected control of the indicated substrates). The data are the mean ± SE for three experiments. (c) TVEC traces of oocytes injected with water only, 5 ng cRNA of AMT1;1 only, or 5 ng AMT1;1 + 0.5 ng CIPK15, and perfused with NH_4_Cl at the indicated concentrations. Similar results were obtained in at least three independent experiments using different batches of oocytes.

**Fig. 2.**
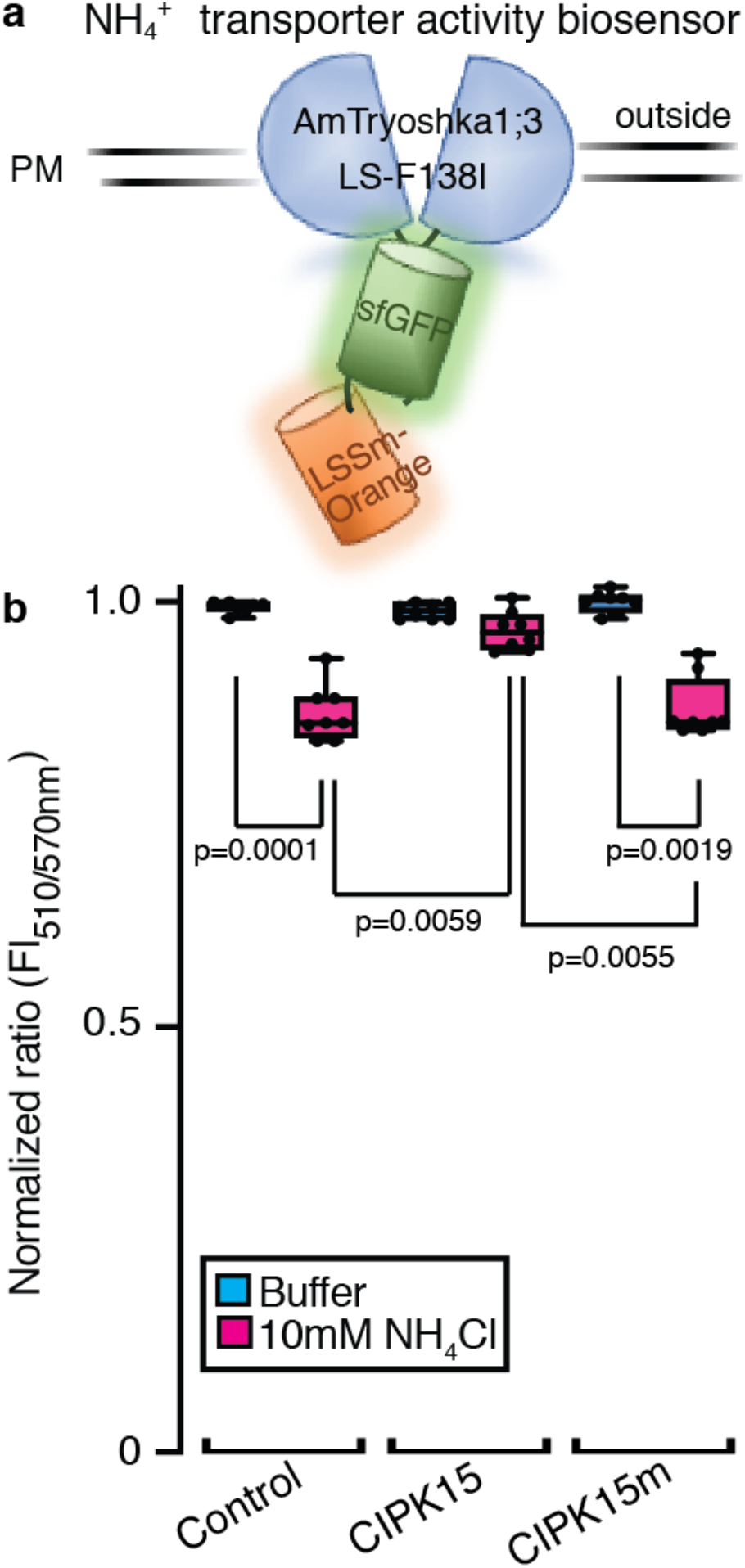
CIPK15 inhibited AmTryoshka1;3 LS-F138I activity in yeast. (**a**) Schematic representation of AmTryoshka1;3 LS-F138I [15]. (**b**) CIPK15 reduced NH_4_^+^-triggered AmTryoshka1;3 LS-F138I [15] responses in yeast. Amtryoshka1;3 LS-F138I was co-expressed with control (vector only), CIPK15 and CIPK15m (inactive mutant). Results of normalized fluorescence ratio (normalized to buffer control=1, *λ* _exc_ 440 nm, ratio= FI_510nm_/_570nm_) after addition of NH_4_Cl as represented by box and whiskers (mean ± SE, n=8). Center lines show the medians; box limits indicate the 25th and 75th percentiles as determined by Prism software; whiskers extend 1.5 times the interquartile range from the 25th and 75th percentiles, outliers are represented by dots. p, significant change as shown in the figure (Two-Way ANOVA followed by Tukey’s post-test). PM = plasma membrane.

### NH_4_^+^-induced *CIPK15* mRNA accumulation

In a screen of different heterologous systems for proteins that can affect AMT activity we identified CIPK15 from Arabidopsis as a negative regulator. We therefore tested whether *CIPK15* may be regulated at the transcriptional level or regulate the AMT activity by NH_4_^+^. To test whether CIPK15 may be linked to NH_4_^+^ nutrition, the expression of CIPK15 in wild-type root in response to addition of 1 mM NH_4_^+^ was examined. Similar to *AMT1;1*, *CIPK15* mRNA levels increased by about 10-fold less than one hour after adding NH_4_^+^ (Fig. 3). These data indicate *CIPK15* is NH_4_^+^ inducible and plays roles in response to NH_4_^+^ nutrition.

**Fig. 3.**
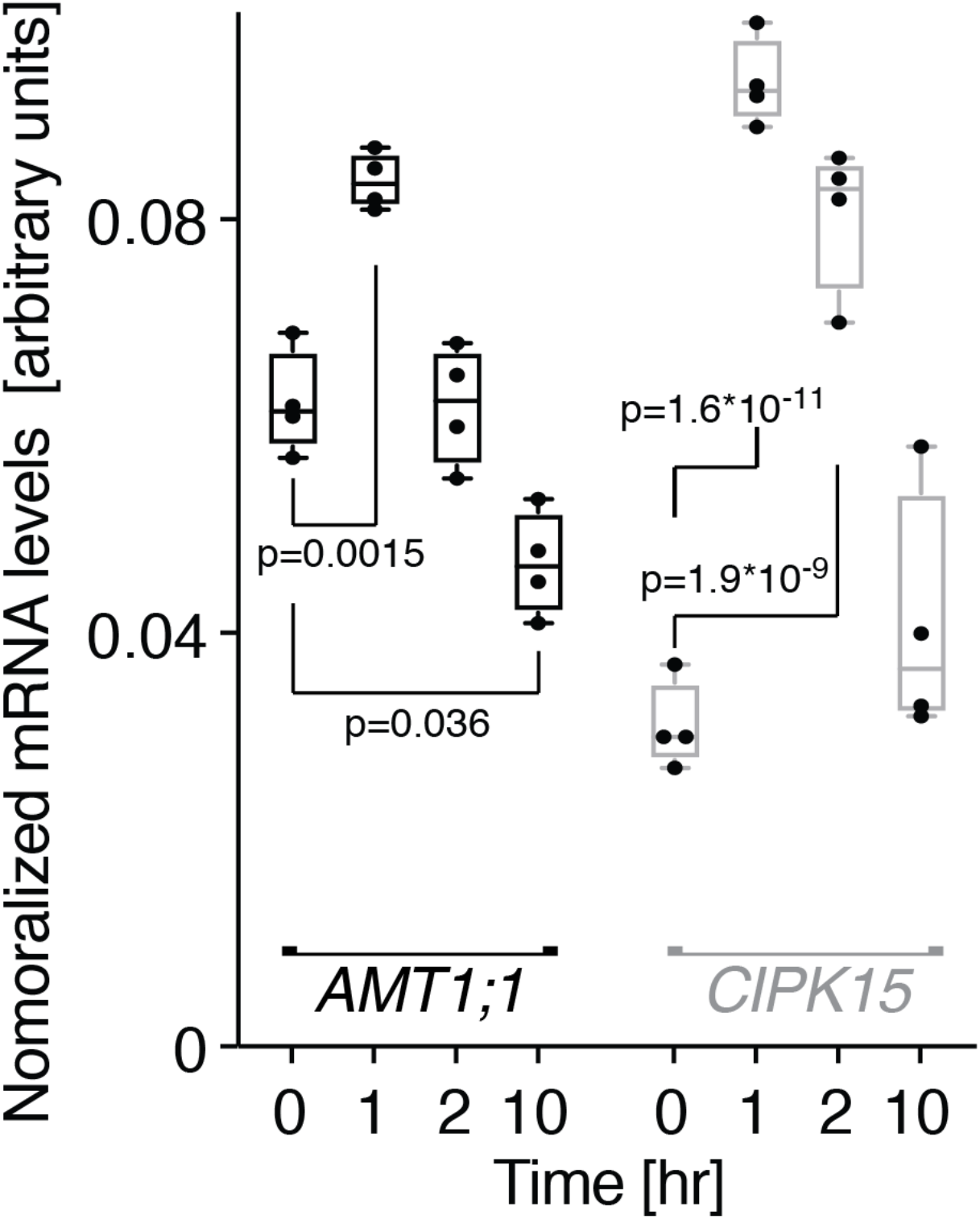
NH4^+^-triggered *CIPK15* mRNA accumulation. qRT-PCR analyses of *AMT1;1* and *CIPK15* mRNA levels in roots after over 10 h after addition of 1 mM NH_4_^+^. Levels were normalized to *UBQ10* [mean ± SE for four independent experiments (each experiment n >50, total n >200)]. p, significant change for mRNA levels of *AMT1;1* and *CIPK15* at 1, 2, and 10 h compared to at 0 h (Two-Way ANOVA followed by Tukey’s post-test).

### CIPK15 can interact with different AMTs

To test whether CIPK15 can interact with AMT1 isoforms, yeast split-ubiquitin interaction growth and β-galactosidase staining and filter assays were used [36]. AMTs (including 1;1, 1;2, 1;3, and 1;5) were fused to Cub (N-terminal ubiquitin domain fused to the artificial protease A-LexA-VP16 (PLV) transcription factor) and CIPK15 to NubG (N-terminal ubiquitin domain Ile-13 (NubI, positive control)) replaced by Gly; reduced affinity for Cub). Plasmids expressing AMTs, CIPK15, and controls (NubI and G) were expressed in yeast. Qualitative and quantitative assays (yeast interaction growth and β-galactosidase assays) demonstrated that CIPK15, but not CIPK19, can interact with AMT1;1 and several different AMTs (Figs. 4a-c, Fig S6, control for Fig 4a, and Fig. S7). The specificity of CIPK15-AMT1;1 interaction was further supported by split-fluorescent protein interaction assays in which different combinations of reconstitution of YFP fluorescence from AMT1;1 and CIPK15 or CIPK19 were used in *N. benthamiana* leaves (Fig. 4d and Fig. S8). Together, the protein interaction results *in vitro* and *in vivo* indicate that CIPK15 can interact specifically with several members of the AMT1 family.

**Fig. 4.**
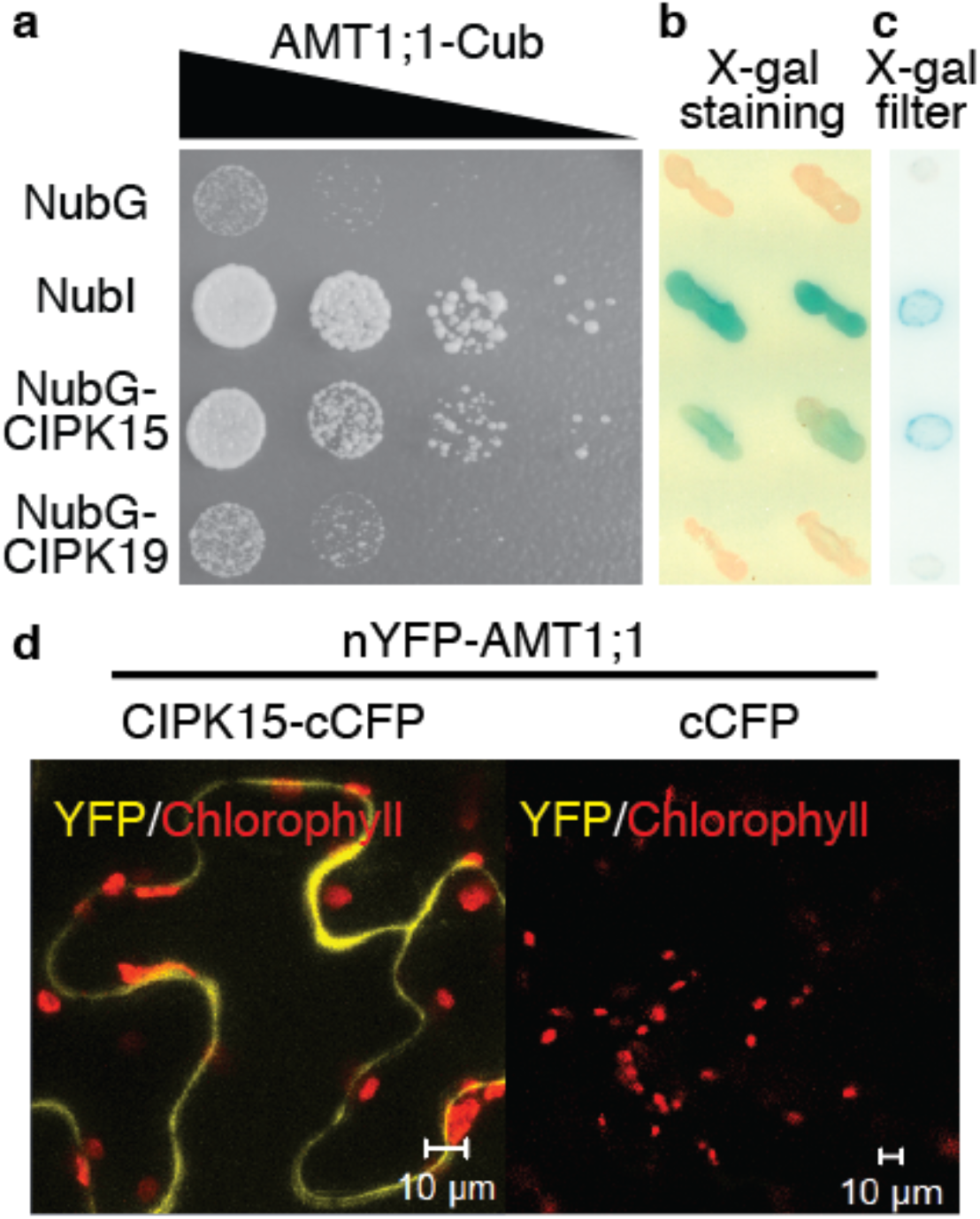
CIPK15 can interact with AMT1;1. Interaction growth assay (**a**), β-galactosidase staining (**b**) and filter (**c**) assay in yeast; and split-fluorescent protein interaction assay (**d**) in tobacco leaves for AMT1.1 and CIPK15 protein interactions. (**a**) Plasmids expressing AMT1;1 and CIPKs were expressed in yeast. Interaction indicated by growth on SD-Trp -Leu -His. Growth on SD-LT as control (Fig. S5). Comparable results were obtained in three independent experiments. (**b-c**) Interaction of CIPKs and AMT1;1 in a split-ubiquitin system detected by X-Gal staining and filter assays using full-length AMT1;1-Cub-PLV as bait and NubG, NubI, and NubG-full-length CIPKs as prey. NubI/NubG served as positive (blue color) and negative controls, respectively. (**d**) Split-fluorescent protein interaction assay for AMT1;1 and CIPK15. YFP/chlorophyll, merged image of fluorescence and chloroplast. Reconstitution of YFP fluorescence from nYFP-AMT1;1 + CIPK15-cCFP and nYPF-AMT1;1 + cCFP (negative control). Comparable results with different combinations shown in Fig. S7.

### CIPK15 is necessary for NH_4_^+^-triggered phosphorylation of T460 in AMT1

AMTs contain multiple possible phosphorylation sites [45]. T460 in the conserved CCT, which immediately follows transmembrane spanning domain XI, plays a key role in the allosteric regulation of AMT1;1 [24]. To test whether CIPK15 is necessary for NH_4_^+^-triggered phosphorylation of T460, we identified two *cipk15 knockout* mutants and analyzed the AMT1:1 phosphorylation status (Fig. S9a-b). Growth of the mutants on MS media and in soil did not indicate any obvious phenotypic differences compared with the wild-type (Fig. S9c). Phospho-specific antibodies were used to test for CIPK15-mediated NH_4_^+^-triggered phosphorylation of T460 using protein gel blots. The *knockout* line AMT-*qko* (quadruple amt mutant) combines T-DNA insertions in *AMT1;1*, *1;2*, *1;3*, and *2;1,* was used as a negative control [32]. After 7 days growth on MS media (high NH_4_^+^), AMT1;1 protein levels were not different in the *cipk15* mutants and wild-type; however, the phosphorylation levels of AMT1 in the wild-type and *cipk15* mutants were low (Fig. S10). In the wild-type, phosphorylation of AMT1 increased substantially within 1 hour of exposure of N-starved plants to NH_4_^+^; AMT1 phosphorylation was undetectable in the *cipk15* mutants (Fig. 5). We therefore conclude that *CIPK15* is necessary for NH_4_^+^-triggered phosphorylation of AMT1.

**Fig. 5.**
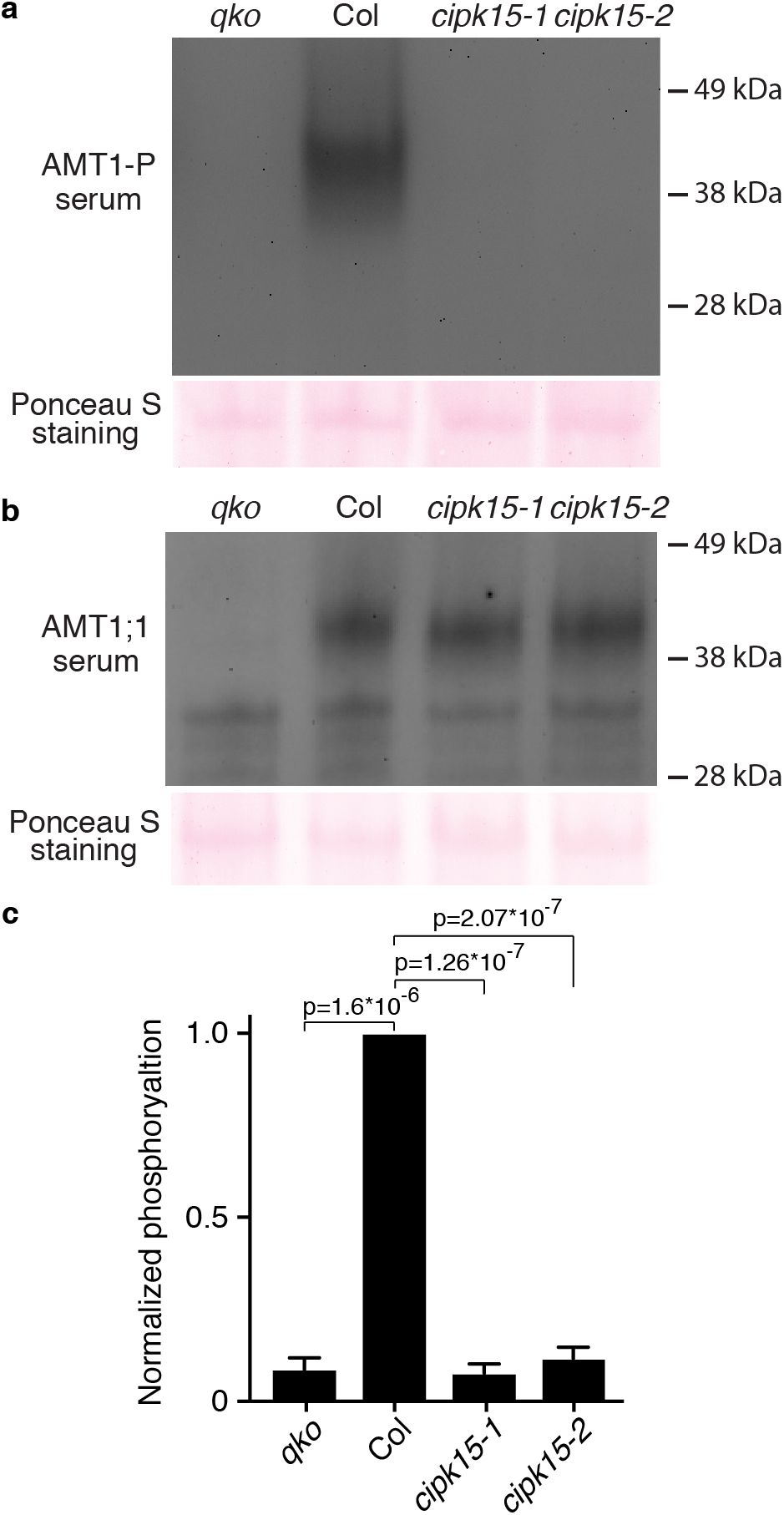
AMT1;1-T460 phosphorylation is reduced in *cipk15* mutant plants. Plant seedlings were germinated and grown for 7 days in half-strength MS medium with 5 mM KNO_3_ as the sole nitrogen source, then starved for 2 days in half-strength MS medium without nitrogen. Seedlings were treated with 1 mM NH_4_Cl for 1 h, membrane fractions were isolated and probed with anti-AMT1-P antibodies (**a**) and anti-AMT1;1 antibodies (**b**) [25]. Ponceau S staining served as a loading control. Quantification of phosphorylation of AMT1-P levels normalized to Ponceau S staining and relative to wild-type shown in (**c**). Corresponding data and replications were obtained in three independent experiments. Data (c) are the mean ± SD for three experiments. p, significant change compared to wild-type as shown in figure (Two-way ANOVA followed by Tukey’s post-test).

### *cipk15* mutant shows high 15NH_4_^+^ uptake activity and NH_4_^+^ accumulation

If CIPK15 is a key regulator that is necessary for T460 phosphorylation, one would predict that *cipk15* mutants should accumulate more NH_4_^+^ and show elevated sensitivity to NH_4_^+^. To determine whether CIPK15 may be able to affect ammonium uptake and NH_4_^+^ toxicity in plants, *cipk15* mutants were exposed to NH_4_^+^. Both *cipk15* mutants were hypersensitive to NH_4_^+^ but not NO_3_^−^. After NH_4_^+^ pretreatment, *cipk15* mutant seedlings accumulated higher amounts of NH_4_^+^ compared to the wild-type (Fig. 6a). Direct analysis of ^15^NH_4_^+^ uptake showed that *cipk15* mutants imported more NH_4_^+^ relative to the wild-type (Fig. 6b). Together, our data showed that CIPK15 is necessary for NH_4_^+^-triggered inhibition of AMT1-mediated NH_4_^+^-uptake.

**Fig. 6.**
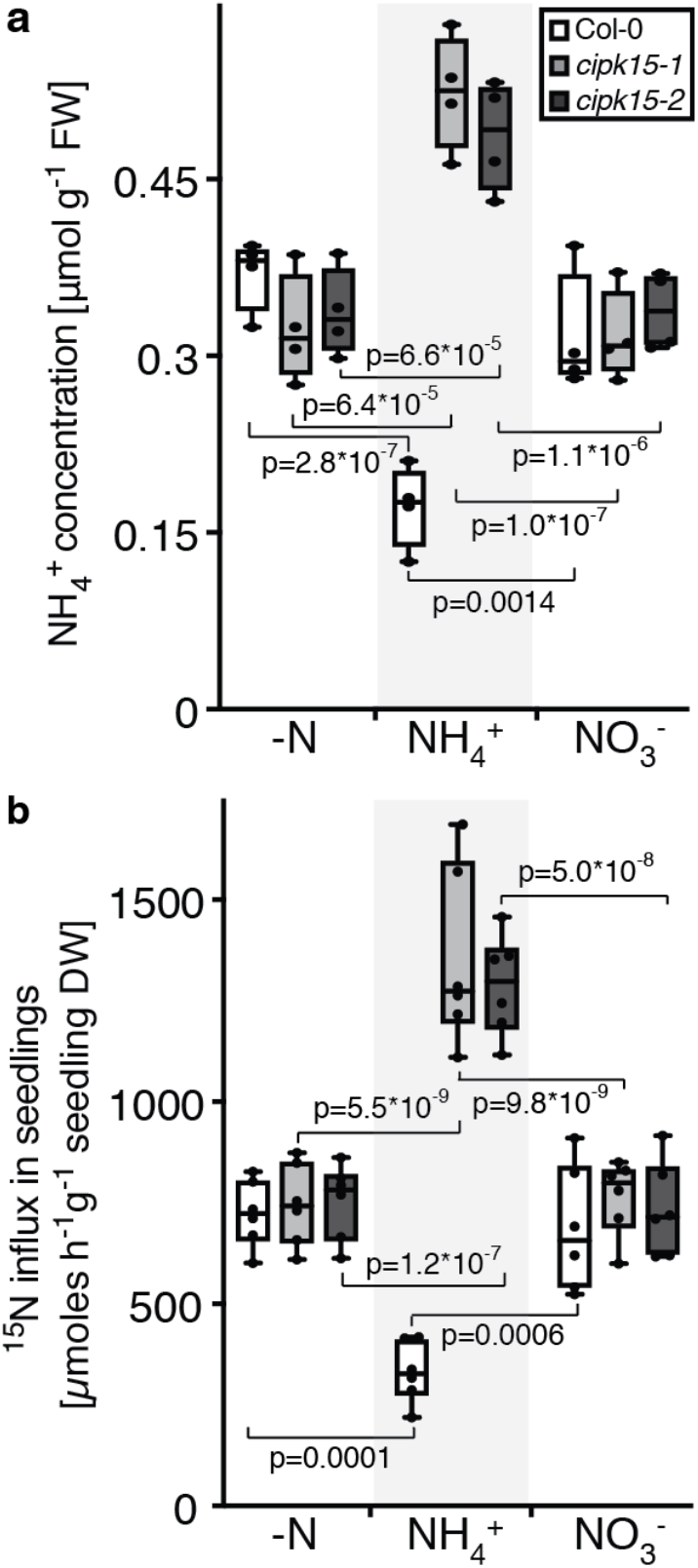
NH4^+^ content and transport in *cipk15* mutants. Plant seedlings were germinated and grown for 7 days in half-strength MS medium with 5 mM KNO_3_ as the sole nitrogen source, then all seedlings were starved for 2 days in half-strength MS medium without nitrogen. For NH_4_^+^ content analyses (**a**) seedlings were collected after being starved for 2 days (-N), or after with 1 mM NH_4_Cl (NH_4_^+^), and 1mM KNO_3_ (NO_3_^−^) for 1 h. For ^15^N labeled uptake, (**b**) seedlings were collected after being starved for 2 days (-N) (1 mM ^15^NH_4_Cl was used for 15 mins for ^15^N-labeling), or after treatment with 1mM NH_4_Cl (NH_4_^+^), and 1mM KNO_3_ (NO_3_^−^) for 1 h (1 mM ^15^NH_4_Cl was used for last 15 mins for N^15-^labeling for conditions of NH_4_^+^ and NO_3_^−^). Each data point represents different experiments, in which seedlings n > 15, total n > 60) in Col-0 and two *cipk15* knockout mutants and presented as box and whiskers. Center lines show the medians; box limits indicate the 25th and 75th percentiles as determined by Prism software; whiskers extend 1.5 times the interquartile range from the 25th and 75th percentiles, outliers are represented by dots. p, significant change compared to before submergence (Two-way ANOVA followed by Tukey’s post-test).

### CIPK15 is a key factor for NH_4_^+^ tolerance in *Arabidopsis*

High levels of NH_4_^+^ negatively impact primary root growth [46–48]. To test whether high accumulation of NH_4_^+^ in *cipk15* mutant and NH_4_^+^-induced phosphorylation of AMT1;1 by CIPK15 affects NH_4_^+^ sensitivity, root growth was analyzed in the presence or absence of NH_4_^+^. Primary root length was not significantly different in the wild-type and *qko* mutants in media containing nitrate as the sole nitrogen source (Fig. S11). By contrast, primary root length of wild-type was dramatically reduced in media containing NH_4_Cl or MeA (Fig. 7 and Fig. S11). Notably, *cipk15* mutants were hypersensitive to NH_4_^+^, but not nitrate, as evidenced by shorter primary root length compared with wild-type and *qko* mutant plants (Fig. 7a and Fig. S12). Ammonium can be taken up via AMTs or K^+^ channels. By contrast, the NH_4_^+^ analog methylammonium (MeA) is transported specifically via AMTs. *cipk15* mutants were also hypersensitive to MeA, further supporting the hypothesis that CIPK15 is necessary for limiting AMT1;1 activity and that the effects observed for NH_4_^+^ can be related directly to the AMTs that contain the conserved domain including T460 (Fig. 7b). CIPK15 has been found to be involved in other processes or interactions with CBL1/4; however, there was no effect on AMT1;1 activity in *Xenopus* oocytes when co-expressed with CBL1 or a constitutively active form of CIPK19 with CIPK15 (Fig. S2), and no effect on primary root length in *cbl4* and *cipk19* mutants when they were exposed to 20 mM NH_4_Cl and KNO_3_ (Fig. S13) indicating that the effects observed with respect to ammonium toxicity are specific. Taken together, we conclude that CIPK15 activity is necessary for limiting NH_4_^+^ uptake by AMT1;1 when roots are exposed to NH_4_^+^ or MeA.

**Fig. 7.**
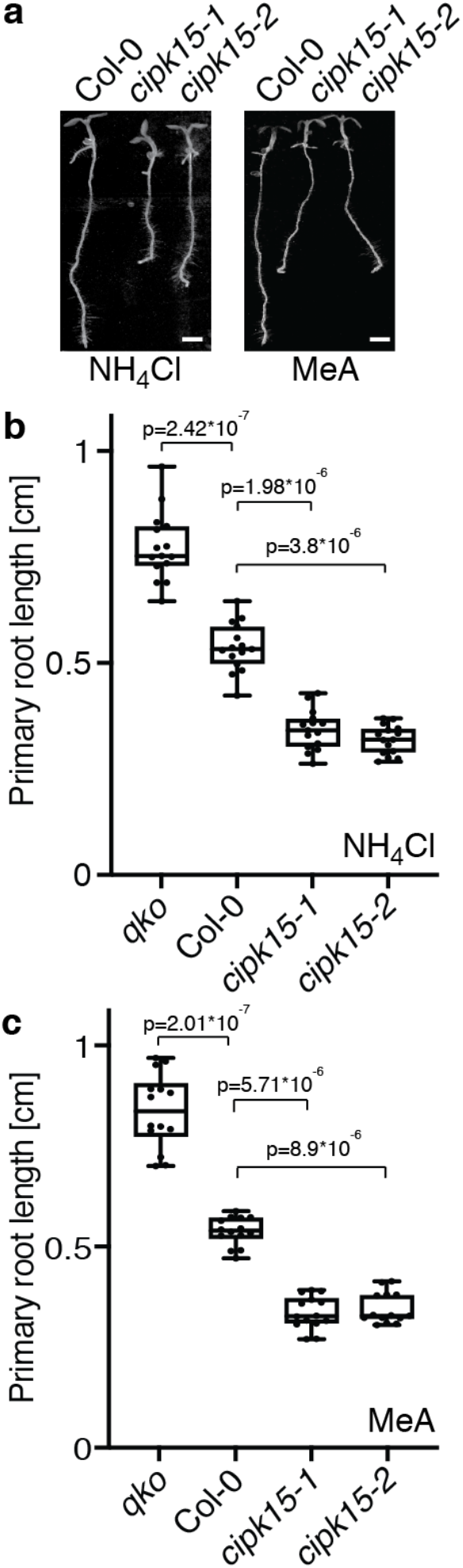
Hypersensitivity of *cipk15* mutant plants to NH4^+^and MeA. Representative images (**a**) and quantification results of primary root length of plants grown on plates containing 20 mM NHCl_4_ (**b**) or 20mM MeA (**c**). Primary root length in wild-type (Col-0), *qko* mutant, and *cipk15* mutants on 20 mM NH_4_Cl (**b**) or on 20 mM MeA (**c**) are presented as box and whiskers. Center lines show the medians; box limits indicate the 25th and 75th percentiles as determined by Prism software; whiskers extend 1.5 times the interquartile range from the 25th and 75th percentiles, outliers are represented by dots (means ± SE; n ≥15). p, significant change of *qko* mutant and *cipk15* mutants compared to wild-type plants (Two-way ANOVA followed by Tukey’s post-test). Scale bar: 0.1cm.

## DISCUSSION

Here, we identified the protein kinase CIPK15 as a key component in the NH_4_^+^-induced downregulation of ammonium uptake in Arabidopsis. CIPK15-mediated allosteric regulation of AMT1 activity may explain the observation that under field conditions, NH_4_^+^-uptake activity is negatively correlated with the external concentration of NH_4_^+^ concentrations in the soil [49].

### Ammonium toxicity

Most plants are sensitive to high levels of NH_4_^+^ and supply with NH_4_^+^ alone typically causes symptoms of growth retardation [16]. Animals and fungi are sensitive to NH_4_^+^ as well, and recent work demonstrates that bacteria are also sensitive to NH_4_^+^. It is thus not surprising that ammonium uptake is under strict control and that the uptake rate is negatively correlated with the history of ammonium exposure [22]. Key questions are how toxic levels can be prevented, how the regulatory networks operate that limit ammonium accumulation and how and where the cells sense ammonium, intracellularly or at the cell surface. The extreme conservation of the CCT in AMTs across kingdoms, even in cyanobacteria and archaebacteria as well as the dominant nature of mutations in the yeast homolog MEP1 piqued our interest and led to studies of the role of the CCT in AMT regulation [24, 50]. Genetic, biochemical and structural analyses have demonstrated that AMTs are triple-barreled transporters that are allosterically regulated. Regulation is mediated by the CCT, which interacts with the respective neighboring subunits for transactivation [24]. A conserved residue, T460 in AMT1;1, T472 in AMT1;2, and T464 in AMT1;3 is phosphorylated in response to addition of NH_4_^+^ [51, 52]. We therefore hypothesized that either a receptor-like kinase or a cytosolic kinase is required for the feedback inhibition.

### CIPK15 is necessary and sufficient for feedback inhibition

CIPKs are known to be involved in the regulation of the activity of diverse sets of transporters including AKT1, SOS1, NPF6.3, IRT1, etc., we therefore hypothesized that specific members of the CIPK family might be able to phosphorylate AMT1;1. To accelerate the screen, we co-expressed sets of five CIPK genes together with AMT1;1 and monitored AMT activity using TEVC. Based on our functional interaction screen assays, we identified and deconvoluted one of the mixtures that led to reduced AMT1;1 activity. CIPK15 by itself was sufficient to substantially inhibit AMT1;1 activity. The inhibition effect on AMT1;1 activity was still obtained when lower amounts of *CIPK15* cRNA were co-injected with *AMT1;1* in oocytes, and CBL1 did not cause the activation of AMT1;1 activity in oocyte. We cannot exclude the possibility that some AMT1;1 activity remains even in the inhibited state in the oocyte system, but the activity was below the detection limit. Commercial oocytes are often lower quality compared to oocytes isolated freshly from locally held frogs, thus it is conceivable that experiments in which higher AMT activity can be detected CIPK15 may also reveal remaining AMT1 activity. However, our data demonstrate that, when co-expressed with AMT1;1 in oocytes, CIPK15 inhibits AMT1;1 activity. Moreover, upon functional interaction assay in yeast, CIPK15 also inhibited NH_4_^+^-induced fluorescence change in the transport activity biosensor AmTryoshka1;3, indicating that CIPK15 can affect the activity of multiple AMT paralogs. The effect of CIPK15 on the AMTs is likely direct and specific, since CIPK15, but not CIPK19 can interact with AMT1;1 or multiple AMTs and tune AMT activities. Importantly, mutant analyses demonstrate that CIPK15 is also necessary for NH_4_^+^-triggered AMT1;1 phosphorylation (T460). *cipk15* mutants took up and accumulated more NH_4_^+^, and were hypersensitive to NH_4_^+^ and the analog MeA. The MeA sensitivity of *cipk15* mutants intimate that the effects observed with respect to ammonium toxicity are due to inhibition of AMT activity, since MeA is transported by AMTs but not by potassium channels. Since CIPK15 is a factor produced in the cytosol, the action of the kinase is intracellular. This work, therefore, identifies the key kinase for AMT regulation, which represents a major step forward for the elucidation of the full regulatory circuit. AMT1;2 also plays an important role in NH_4_^+^ uptake. Data from other groups may indicate that in oocytes AMT1;2 mediates larger ammonium-induced inward currents when compared to AMT1;1 [53]. It remains open, whether the larger currents are due to different quality of oocytes from in house versus commercial facilities. The next experiments will need to address where and how NH_4_^+^ is sensed. CIPK15 and the AMTs may be useful tools to unravel the remaining steps in the regulatory circuitry.

### The relative role of CIPK15 and CIPK23 in AMT regulation

Recent work has indicated that another CIPK, namely CIPK23 plays a role in the regulation of AMT1;1 and AMT1;2 [53]. The authors showed that CIPK23 can interact with AMT1;1 and AMT1;2 but did not observe an interaction with AMT1;3. Here, we identified an interaction between CIPK15 and AMT1;1 by using split-ubiquitin yeast two-hybrid assays in yeast, split-fluorescent protein interaction assays in *N. benthamiana* leaves, and functional interaction by TEVC in *Xenopus* oocytes. Interactions of CIPK15 with AMT1;2 and AMT1;3 were also identified by using split-ubiquitin yeast two-hybrid in yeast, and a functional interaction of CIPK15 with AMT1;3 was validated with the help of a ratiometric NH_4_^+^ transporter activity reporters in yeast. Consistent with the conservation of the domain surrounding the phosphorylation site (T460 in AMT1;1), the protein interaction and functional assay results indicate that CIPK15 likely affects activity of all three AMT1 paralogs. According to public transcriptome databases (e.g. TAIR, Genevestigator), *CIPK23* and *CIPK15* appear to be expressed in AMT1;1-expressing tissues. Here, we also found that *CIPK15* and *CIPK23* mRNA increased in response to NH_4_^+^ addition (Fig. S14). Notably, *CIPK15* mRNA accumulation triggered by 1mM NH_4_^+^ was about three and half-fold higher relative to the *CIPK23* mRNA accumulation, absolute levels of *CIPK15* are similar after ammonium addition compared to *CIPK23*. Consistent with the interaction, coexpression of CIPK23 in the presence of CBL1 led to about a two-fold lower current for AMT1;2 in *Xenopus* oocytes, while CIPK15 led to essentially complete loss of detectable AMT1;1-mediated currents. The data from the two labs are not directly comparable, since Straub et al. observed larger currents when analyzing the effect of CIPK23 on AMT1;2. AMT1;3 activity was also impaired by CIPK15 as shown using AmTryoshka1;3. While the experiments were not performed side by side, these data may indicate that CIPK23 plays a different and less prominent role as compared to CIPK15. In *cipk23* mutants, AMT1;1-GFP phosphorylation was reduced by ~20% for AMT1;1 and ~40% for AMT1;2. In comparison, *cipk15* mutants completely lost detectable AMT phosphorylation. In *cipk23* mutants, the shoot dry weight was reduced, and they showed higher ^15^NH_4_^+^ uptake in the presence of ammonium relative to the wild-type. However, *cipk23* mutants displayed no difference regarding hypocotyl length when exposed to 20 mM ^15^NH_4_^+^. Well characterized CIPK15-interactors, CBLs, did not have an apparent effect on AMT1;1 activity in *Xenopus* oocytes, nor was NH_4_^+^ toxicity in *cbl* mutants affected. Taken together the data indicate that multiple CIPKs can affect AMT activity with different efficacy, possibly different tissue specificity, different specificity for AMT paralogs, and with differing dependence on CBLs.

Taking the results together, this work identified a key component in the NH_4_^+^ feedback inhibition network, namely the protein kinase CIPK15, which directly interacts with AMTs to phosphorylate the conserved threonine in their C-terminus to adjust ammonium uptake and dependence on the external NH_4_^+^ concentration. The remaining open questions in the field are how the plant senses the ammonium concentration and how it activates CIPK15. CIPK15 has been reported involved in the ABA signaling pathway and phosphorylation of ERF7, an APETALA2/ EREBP-type transcription factor [54, 55], and more recently, CPK32 was shown to play a role in the regulation of AMT activity, it will be interesting to explore the ABA response and the interrelationship between CPK32 phosphorylation of residues downstream of T460 and the two CIPKs [56].

## ACKNOWLEDGEMENTs

We would like to thank the Dr. Jörg Kudla (U. Münster, Germany) for providing plasmids containing *CIPK* ORFs for oocytes experiments. We thank Prof. Charles Brearley, School of Biological Sciences, and Dr. Sarah Wexler, Science Analytical Facility, (U. East Anglia, Norwich Research Park) for N^15^ analyses. We thank Academia Sinica Advanced Optics Microscope Core Facility for technical support for fluorescence imaging. The core facility is funded by Academia Sinica Core Facility and Innovative Instrument Project (AS-CFII-108-116). We thank Anita K. Snyder and Miranda Loney for English editing. This research was supported by Academia Sinica, Taiwan, and Ministry of Science and Technology, Taiwan, Grants MOST 105-2311-B-001-045 and 106-2311-B-001-037-MY3 (C.-H.H.) and Deutsche Forschungsgemeinschaft (DFG, German Research Foundation) under Germany’s Excellence Strategy – EXC-2048/1 – project ID 390686111, SFB 1208 – Project-ID 267205415, as well as the Alexander von Humboldt Professorship (WBF).

## AUTHOR CONTRIBUTIONS

Conceptualization, C.-H.H. and W.B.F.; Methodology, C.-H.H., H.-Y.C., Y.-N.C., and H.-Y.W.; Investigation, C.-H.H., H.-Y.C., Y.-N.C., H.-Y.W., and Z.-T.L.; Writing, C.-H.H. and W.B.F.; Supervision, C.-H.H. The work was initiated by C.-H.H. in W.B.F.’s lab. The major parts were performed in C.-H.H. own lab.

